# The telencephalon is a neuronal substrate for systemic inflammatory responses in teleosts via polyamine metabolism

**DOI:** 10.1101/2024.03.03.583242

**Authors:** Amir Mani, Farah Haddad, Daniel R. Barreda, Irene Salinas

**Author notes:** Irene Salinas **Email:**. **Author Contributions:** Conceptualization: IS, AM; Investigation: AM, FH, DB, IS; Methodology: AM, IS; Formal analysis: AM, FH; Writing-original draft: AM, IS; Writing-reviewing and editing: AM, FH, DB, IS. **Competing Interest Statement:** Authors have no competing interests. **Classification:** Biological sciences. Immunology and Inflammation.

## Abstract

Systemic inflammation elicits sickness behaviors and fever by engaging a complex neuronal circuitry that begins in the preoptic area of the hypothalamus. Ectotherms such as teleost fish display sickness behaviors in response to infection or inflammation, seeking warmer temperatures to enhance survival via behavioral fever responses. To date, the hypothalamus is the only brain region implicated in sickness behaviors and behavioral fever in teleosts. Yet, the complexity of neurobehavioral manifestations underlying sickness responses in teleosts suggests engagement of higher processing areas of the brain. Using two *in vivo* models of systemic inflammation in rainbow trout, we find canonical pyrogenic cytokine responses in the hypothalamus whereas in the telencephalon *il-1b* and *tnfa* expression is decoupled from *il-6* expression. Polyamine metabolism changes, characterized by accumulation of putrescine and decreases in spermine and spermidine, are recorded in the telencephalon but not hypothalamus upon systemic injection of bacteria. While systemic inflammation causes canonical behavioral fever in trout, blockade of bacterial polyamine metabolism prior to injection abrogates behavioral fever, polyamine responses and telencephalic but not hypothalamic cytokine responses. Combined, our work identifies the telencephalon as a neuronal substrate for brain responses to systemic inflammation in teleosts and uncovers the role of polyamines as critical chemical mediators in sickness behaviors.

**Significance Statement:** Systemic inflammation induces neuroimmune responses in the brain resulting in sickness behaviors and fever. In endotherms, sickness behaviors and fever are initiated in the hypothalamus but also engage a complex neuronal circuitry in higher areas of the brain. In ectotherms, only the hypothalamus has been linked to sickness behaviors and behavioral fever. Here we demonstrate that the telencephalon, a critical region of the teleost brain responsible for higher order processing, mounts pyrogenic cytokine responses to systemic inflammation in teleosts that are different from those of the hypothalamus. We identify polyamine metabolism as a core response of the teleost telencephalon to systemic inflammation and report that bacterial polyamines are triggers of behavioral fever in rainbow trout. Our work uncovers a previously unrecognized role for the telencephalon and polyamine metabolism in sickness behaviors and behavioral fever in teleosts with implications in fish health and fish conservation.

## Introduction

The bidirectional communication between the periphery and the brain regulates homeostasis and disease states in animals (1–8). Systemic inflammation results in sickness behaviors, such as anorexia, depression, lethargy, sleepiness, social withdrawal, and sometimes fever responses. Sickness behaviors and fever are cytokine orchestrated, complex neurobehavioral phenomena that involve multiple neural substrates (9–13). In mammals, the Median Preoptic Nucleus (MnPO) of the hypothalamus is the first region to detect systemic inflammatory signals (11–14) but extensive experimental evidence indicates that the hypothalamus is not the only brain region responding to systemic inflammation. For instance, systemic inflammation engages neurons in the mouse insular cortex and these neurons can evoke past inflammatory states in the periphery following optogenetic neuronal activation (15, 16). Furthermore, bacterial lipopolysaccharide (LPS) injection, the gold standard model for inducing sickness behaviors in mammals, results in significant neuronal activation not only in the hypothalamus but also in key limbic regions, including the amygdala, hippocampus, nucleus accumbens, and the stria terminalis, indicating a broad neuroanatomical network responding to systemic inflammation (17, 18). Similarly, in rats, LPS injection triggers astrocyte activation within the striatum and hippocampus with significant volumetric expansions occurring in the splenium, retrosplenial cortex, and pericallosal hippocampal cortex at 2 hours post-injection (hpi) (19). Combined, mammalian studies indicate that systemic inflammation disturbs several higher-order cortical areas (13, 15, 20, 21), their connectivity across the entire brain yet to be fully resolved.

Ectotherms, like mammals, also display sickness behaviors in response to infection, stress and inflammation (22–26). For instance, teleosts become lethargic, stop feeding and seek warmer water temperatures in a process known as behavioral fever (27–32). This behaviorally induced fever provides a survival advantage, as it enhances the immune system’s efficacy, hinders the proliferation of pathogens and resolves wound healing more effectively (27, 28, 31–33). Similar to mammals, systemic and brain increases in pyrogenic cytokines like IL1β, TNFα, and IL6, alongside PGE2, have been identified as key players in teleosts behavioral fever responses. Furthermore, the preoptic area of the hypothalamus (POA) is thought to be, as in mammals, the main brain region responding to systemic infection and driving behavioral fever responses in ectotherms (30, 33–35). Yet, expression of the prostaglandin receptor EP3 was detected in the telencephalon and optic lobe of Atlantic salmon (29), suggesting that other brain regions detect systemic inflammation in teleosts. Given the complexity of sickness behaviors and fever in teleosts, we hypothesized that the teleost brain must not only sense and produce proinflammatory cytokines in response to systemic inflammation, but also engage higher order processing areas responsible for decision-making and voluntary movement.

The teleost brain, though structurally different from that of mammals, performs similarly complex functions (36). The telencephalon of teleosts, comprising the pallium and subpallium, is increasingly recognized for its role in higher-order processing including spatial cognition, sensation, and voluntary movement, all of which are integral to behavioral fever (37–43). For instance, the teleost telencephalon plays pivotal roles in spatial navigation (40, 41, 43), avoidance behaviors (38), and the coordination of movement towards warmer environments (41, 44, 45). Interestingly, the dorsal, medial, and ventrodorsal telencephalon in teleosts are analogous to the mammalian hippocampus, amygdala, and basal ganglia, respectively (37–43, 46), all of which respond to systemic inflammation in mammals. Based on these premises, along with the EP3 expression in the teleost telencephalon, we hypothesized that this region would be a likely neuronal substrate candidate in systemic inflammation and behavioral fever responses in teleosts.

Herein we investigate the neuroimmune responses of the rainbow trout brain to systemic inflammation. By comparing the telencephalon to the hypothalamus, we identify sharp differences in how these two brain regions respond to inflammation. We also provide experimental evidence that shifts in polyamine metabolism occur specifically in the telencephalon in response to systemic inflammation, and that bacterial derived polyamines regulate the onset of behavioral fever. Our study reveals a novel neuronal substrate beyond the POA that governs responses to systemic inflammation in higher processing areas of the teleost brain.

## Results

### Systemic microbiota and LPS induce distinct cytokine responses in the trout telencephalon and hypothalamus

To investigate brain neuroimmune responses to systemic inflammation we developed two *in vivo* models in juvenile rainbow trout (Fig. 1A). In the first model, we collected bacteria from the gut of healthy rainbow trout and injected it intravenously (i.v) (fig. S1, A and B). Since ectotherms are highly refractory to *E. coli* LPS injection (47–49), we decided to isolate LPS from the native trout gut microbiota instead of using commercial sources. We first measured LPS levels in the serum of trout injected i.v with the isolated bacteria (Fig. 1B). Notably, LPS levels were only elevated in the blood serum of fish injected with gut bacteria at 4 hpi, returning to baseline levels by 1 day (Fig. 1B). Next, we adjusted how much of the commensal LPS we would need to inject to match this value in both models (Fig. 1A and B). In both models, we noted canonical reductions in fish locomotion, but no mortalities occurred during any of the experiments.

**Figure 1:**
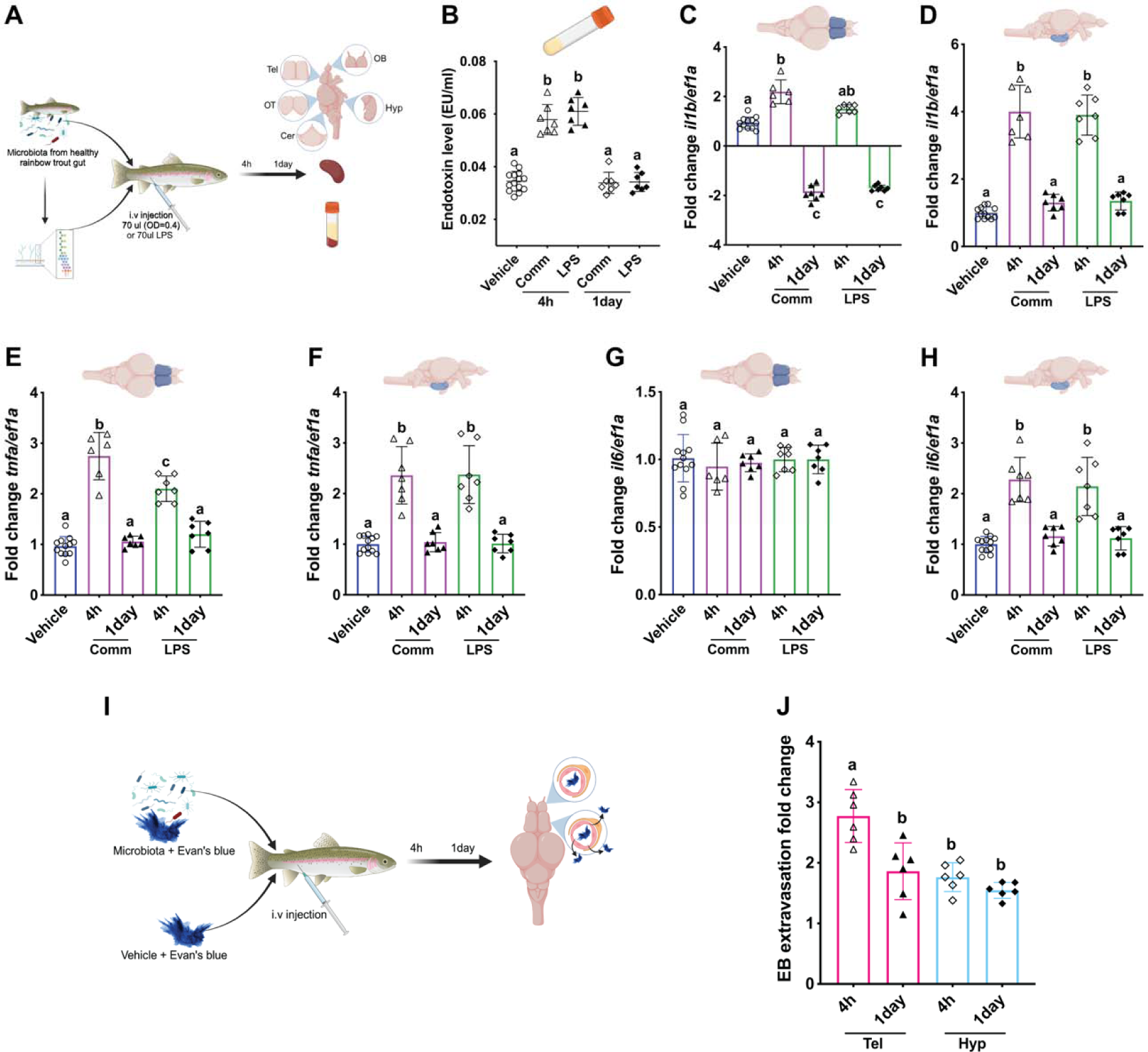
Differential responses in the trout telencephalon and the hypothlamus to systemic inflammation. (A) Schematic representation of the *in vivo* acute commensal translocation models used in the present study. Rainbow trout received an i.v injection of gut commensal bacteria (Comm), isolated lipopolysaccharides (LPS), or vehicle (control). Animals were sampled 4 hours or 24 hours later and blood, spleen and five regions of the brain were sampled for subsequent analysis. (B) Changes in serum endotoxin levels post-injection with gut commensals or LPS (N= 6). Mean fold-change in expression levels of the pro-inflammatory cytokine genes *il-1b*, *tnfa* and *il-6* in the Tel (C, E, G) and Hyp (D, F, H) after injection of gut commensals or LPS compared to vehicle controls as measured by RT-qPCR (N= 6). Vehicles show N=12, N=6 for 4 hours and N=6 for 1 day, with differences between each group. (I) Schematic representation of the *in vivo* Evan’s blue extravasation assay to assess the permeability of the blood-brain barrier in the different experimental groups and brain regions. (J) Analysis of Evan’s blue dye extravasation in the Tel and Hyp, indicating changes in blood-brain barrier integrity post-injection of gut commensal bacteria (N=6). Statistical significance is indicated by differing letters based on one-way ANOVA followed by Tukey’s post hoc test (p<0.05).

Previous studies have established that like mammals, the teleost hypothalamus (Hyp) expresses pyrogenic cytokines in response to systemic or systemic infection (28, 32, 35). However, whether other regions of the teleost brain respond to systemic inflammation is currently unknown. We hypothesized, given the role of the teleost telencephalon (Tel) in higher order processing, that this region would be part of the neuronal circuitry responding to systemic insults. Thus, we compared the expression of pyrogenic cytokine genes in the Hyp and Tel in both of our *in vivo* models. As previously reported in the Hyp, *il-1b*, *tnfa*, and *il-6* expression was significantly higher (2-4-fold increase) in commensal-injected and LPS-injected trout 4 hpi (Fig. 1D, F, and H). Expression levels of all the cytokines returned to baseline levels by 1 day for both the commensal and LPS-injected groups (Fig. 1D, F, and H). In the Tel, *il-1b* and *tnfa* expression rose 2-3 folds at 4 hpi for both commensals or LPS injection (Fig. 1C and E). However, *il-6* expression remained unchanged at all tested timepoints and treatments (Fig. 1G). Like the Hyp, Tel cytokine expression levels returned to basal levels by 1 day. These results indicate that the trout Tel responds to systemic inflammation with a unique cytokine signature compared to the Hyp.

LPS and other systemic inflammation mediators cause breakdown of the blood brain barrier (BBB) in mammals (50, 51). We measured the integrity of the trout BBB in response to the injection of gut-derived bacteria using Evans blue extravasation, a widely used technique to measure BBB permeability (52–54) (Fig. 1I and J). We measured Evans blue extravasation in five different brain regions including olfactory bulb (OB), Tel, Hyp, optic tectum (OT) and cerebellum (Cer). BBB permeability increased by 2-fold compared to vehicle controls in all regions examined by 4 hours (fig. S1, C to E) except for the Tel, where we observed a >3-fold increase in Evans blue extravasation at 4 hpi when compared to baseline levels (Fig. 1J). This increased permeability was transient, with levels decreasing towards the baseline within 1 day post-injection, indicating a temporary disruption of the BBB in the Tel (Fig. 1J). In contrast, other brain regions including the Hyp, showed no significant BBB disruption by 1 day (Fig. 1J and fig. S1, C to E). These data indicate that systemic inflammation preferentially breaks down the Tel BBB compared to other brain regions in rainbow trout.

### Systemic bacteria transiently and selectively penetrate the trout telencephalon

Our two *in vivo* models indicated that the Tel and the Hyp respond differently to systemic inflammation. We have recently reported that salmonids have a resident bacterial community in their brain at steady state (55). We therefore wondered if upon systemic injection of microbiota, bacteria would be able to penetrate the brain. We quantified total bacterial loads in the spleen and five regions of the brain 4 hours and 1 day post-injection using 16s rDNA qPCR. In the Tel, a substantial increase in bacterial loads was observed at 4 hpi with gut commensals. Bacterial presence was transient, as levels at 1 day did not differ significantly from baseline measurements (Fig. 2A). Conversely, the bacterial loads in the Hyp as well as all other brain regions and the spleen remained unchanged (Fig. 2B and fig. S2, E to J). As predicted, the administration of LPS did not change bacterial loads in any tissue sampled at either 4 hours or 1 day post-injection (fig. S2, E to J). We next tested a second model of gut bacterial translocation and systemic inflammation (fig. S3, A). We employed the recently developed DSS-induced colitis in rainbow trout (56) and 1 day after the last DSS gavage, we collected the same tissues as before except the Hyp (fig. S3, A). Similar to the i.v injection model, we noted significant increases in serum endotoxin levels in DSS treated compared to vehicle treated controls (fig. S3, B), but bacterial loads were only significantly higher in the Tel of DSS-treated animals compared to controls, with no other significant changes noted in the other brain regions (fig. S3, C to G). Combined, these experiments indicate that transient and region-specific bacterial translocation occurs in the trout brain during systemic inflammation.

**Figure 2:**
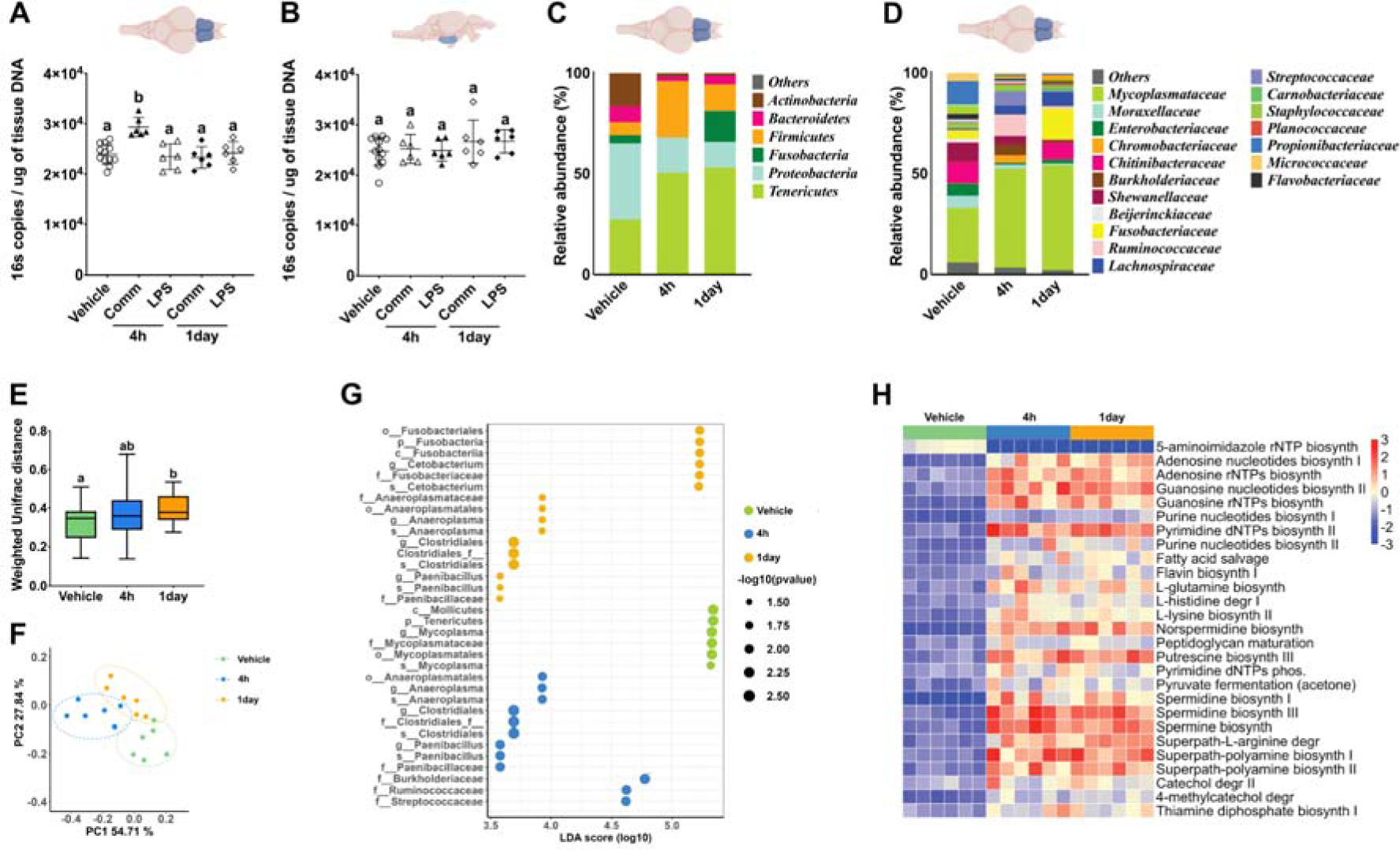
Systemically injected commensals translocate into the telencephalon but not other regions of the trout brain. 16S rRNA gene copies in the Tel (A) and Hyp (B) of rainbow trout at 4 hours and 1 day post bacterial injection (N=6). Vehicles consist of N=12 because they include N=6 vehicle controls for each time point with no differences among them. Statistical significance is indicated by different letters based on Welch’s ANOVA (P<0.05). Relative abundance at the phylum level (C) and family level (D) in the trout Tel 4 hours or 1 day post-injection (N= 6). (E) Weighted UniFrac distances showing significant differences in microbial community structure in the Tel at 4 hours and 1 day post-injection, as determined by the Kruskal-Wallis test (P<0.05) (N=6). (F) Principal coordinate analysis (PCoA) of the Weighted Unifrac distances, with ellipses denoting a 95% confidence interval. (G) Linear discriminant analysis effect size (LEfSe) identifying significantly different taxa in the Tel at specified post-injection time points. (H) Predicted metabolic pathways significantly modified in the microbial communities of the telencephalon 4 hours or 1 day post-injection of commensal bacteria compared to vehicle controls based on PICRUSt2 analysis (P<0.05).

Given that bacteria penetrate the trout Tel, we next sought to evaluate whether shifts in the Tel microbial community occur following i.v bacterial injection using 16S rDNA sequencing. Collected gut bacteria from healthy laboratory rainbow trout were predominantly composed of four bacterial phyla: Tenericutes (72.2%), Proteobacteria (11.2%), Fusobacteria (7.4%), and Firmicutes (7.2%). Further analysis at the family level revealed the bacterial community to consist primarily of eight families: Mycoplasmataceae (72.2%), Fusobacteriaceae (7.2%), Lachnospiraceae (4.9%), Chitinibacteriaceae (3.0%), and Burkholderiaceae (1.9%) (fig. S1, A and B) and were markedly different from the tank water microbial community (fig. S4, A and B). The predominant bacterial phyla identified in the Tel of control animals included Proteobacteria (37.93%), Tenericutes (27.0%), Actinobacteria (16.5%), and Bacteroidetes (7.9%) mainly composed of Mycoplasmataceae (27.04%), propionibacteriaceae (11.3%), Chitinibacteraceae (10.7%) and Shewanellaceae (9.1%) families (Fig. 2C and D). Finally, the bacterial composition in the tank water was distinctly different from that observed in the Tel and composed mostly of members of Acidobacteria (29.5%), Bacteroidetes (31.2%), Proteobacteria (17.3%), and Nitrospirae (6.2%) phyla, such as the families Blastocatellia (uncultured family, 26.6%), Microsillaceae (8.1%), Nitrosospiraceae (6.2%), and Saprospiraceae (11.4%), indicating a unique microbial community compared to that of the Tel (fig. S2, A and B). Following i.v injection of bacteria, there was an increase in the relative abundance of Tenericutes, Firmicutes, and Fusobacteria, whereas Proteobacteria, Bacteroidetes, and Actinobacteria relative abundance decreased compared to the vehicle group at the 4-hour and 1-day time points (Fig. 2C). At the family level, the relative abundance of Mycoplasmataceae, Lachnospiraceae, and Chromobacteriaceae increased at both time points (Fig. 2D). Conversely, the relative abundance of families such as Enterobacteriaceae, Moraxellaceae, and Shewanellaceae diminished at the same intervals (Fig. 2D and G). While the Tel microbial community composition at the 4-hour and 1 day post-injection time points were largely similar, Ruminococcaceae and Streptococcaceae accounted for 10.4% and 7%, respectively, of the overall diversity at 4 hours whereas they were only 1.3% and 0.3%, respectively, at 1 day. At 1 day, Fusobacteriaceae and Chitinibacteraceae relative abundances were 15.4%, and 8.3%, respectively, a much larger percentage than at 4 hours (0% and 0.1%) (Fig. 2D). The Weighted UniFrac distance between the Tel bacterial community of vehicle-injected group and commensal-injected group was significantly different at 1 day (Fig. 2E). Principal coordinate analysis revealed distinct microbial community compositions between the vehicle and commensal injected groups, with the Tel microbial communities of the injected animals remaining closely clustered (Fig. 2F). Alpha diversity metrics (Shannon diversity, Chao1, and Pielou’s evenness) did not show any significant differences in bacteria-injected versus vehicle-injected animals (fig. S4, C to E).

PICRUSt2 analysis revealed shifts in functional pathways of the Tel microbial community following i.v injection of bacteria. A total of 27 pathways were significantly modulated in the Tel microbial community at 4 hour and 1 day compared to vehicle controls. Several polyamine biosynthesis pathways, such as putrescine, spermidine, and spermine biosynthesis were upregulated at both time points compared to controls (Fig. 2H). Additionally, nucleotide biosynthesis, including adenine and guanine pathways, L-glutamine and L-lysine biosynthesis, and degradation pathways like L-histidine degradation and fatty acid salvage were modulated as a result of the bacterial injection treatment (Fig. 2H). These results indicate that, despite bacterial loads dropping by 1 day, the systemic inflammation in the Tel microbial community composition and its predicted functions has longer lasting impacts. Combined, these results show that only specific suites of gut systemic bacteria penetrate the Tel of rainbow trout altering its community composition and predict changes in polyamine metabolism in the Tel bacterial community.

### Systemic inflammation modulates host brain polyamines in a region-specific manner

Polyamines are polycationic organic molecules with two or more amino groups. Polyamines are universal molecules in nature, produced by archaea, bacteria and eukaryotes and are involved in fundamental biological processes such as growth, cell proliferation, differentiation, and death (57–60). Given that the Tel microbial composition showed predicted changes in bacterial polyamine synthesis pathways, we next asked if these shifts had an impact on host polyamine synthesis pathways. Thus, we measure expression levels of four genes encoding for polyamine synthesis enzymes (*odc1*, *odc2*, *samdc1*, and *samdc2*) in the trout Tel and Hyp (Fig. 3A and B). We observed a marked increase in the expression of *odc* genes in the Tel, with two-fold and three-fold elevations at 4 hours and 1 day post-injection of gut commensals, respectively (Fig. 3C and fig. S5, A). Conversely, a significant decrease in the levels of *samdc* transcripts was noted in the Tel, with two-fold and three-fold reductions at 4 hours and 1 day post-injection of gut commensals, respectively (Fig. 3D and fig. S5, C). LPS injection, however, did not cause any changes in *odc* or *samdc* expression in the Tel. Interestingly, *odc* and *samdc* gene expression did not change in the Hyp at any given time point or treatment group (Fig. 3E and F and fig. S5, B and D), indicating the host polyamine responses to systemic bacteria are region-specific.

**Figure 3:**
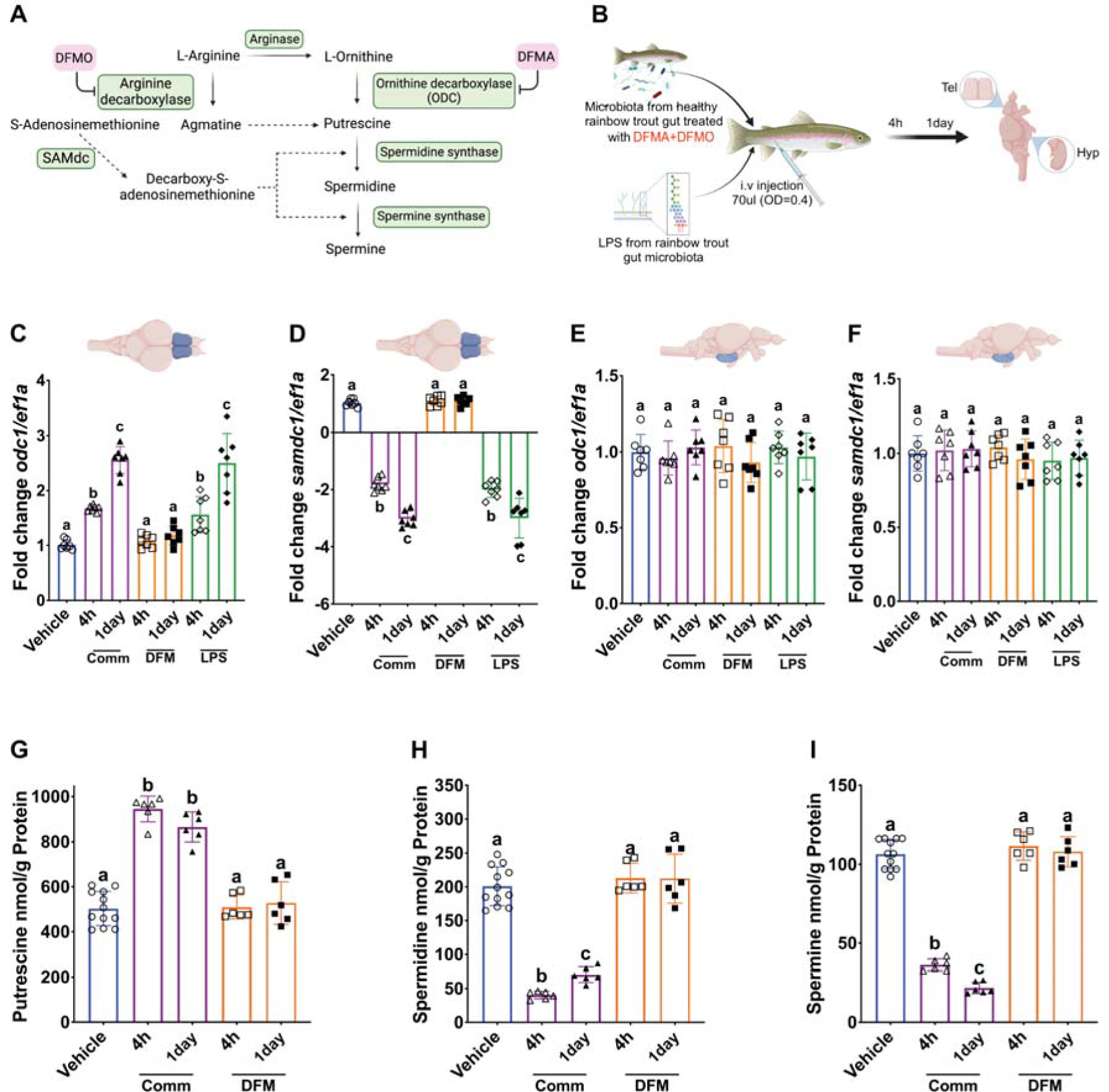
Systemic inflammation results in telencephalic polyamine responses. (A) Diagram depicting the bacterial polyamine synthesis pathway, with the inhibitory effects of DFMA and DFMO on bacterial ornithine decarboxylase and arginine decarboxylase, respectively. (B) Schematic diagram showing the experimental systemic inflammation model of the study consisting of a single i.v injection of gut commensal bacteria (Comm), gut commensal bacteria treated with DFMA-DFMO (DFM), isolated LPS (LPS), or vehicle controls. Changes in the expression levels of polyamine synthesis enzymes *odc1* and *samdc1* in the Tel (C and D) and Hyp (E and F) post-injection measured by RT-qPCR. The expression levels are reported as the mean fold change compared to vehicle controls (N=7 fish/group). (G-I) Mean concentration of putrescine, spermidine and spermine in the trout Tel 4 hours or 1 day post-injection for vehicle, commensal-injected (Comm) and commensal treated with DFMA-DFMO (DFM) (N=6) fish/group). The vehicle group includes N=12 because it combines N=6 vehicle controls for each time point with no differences among them. Different letters indicate statistically significant groups at 95% interval using one-way ANOVA followed by Tukey’s post-hoc test (p<0.05).

In order to test if host changes in polyamine synthesis gene expression are dependent on the ability of bacteria to make polyamines, we blocked polyamine metabolism in the microbiota prior to injecting them into the host. We treated isolated gut commensals with the polyamine pathway inhibitors DFMA (α-difluoromethylarginine), an irreversible inhibitor of arginine decarboxylase, and DFMO (α-difluoromethylornithine), an irreversible suicide inhibitor of ornithine decarboxylase (fig. S2, A to D). Since polyamines can be essential for bacterial growth (61–64) we first performed *in vitro* experiments to establish the highest dose of DFMA and DFMO that would not compromise bacterial growth (fig. S2, A to D). Based on these assays we selected 0.25 mM for each drug for all subsequent *in vivo* experiments (Fig. 3B). Bacteria were washed three times prior to injection to ensure the removal of DFMA and DFMO (65). We first assessed if treated bacteria were still capable of translocating into the trout Tel and found that polyamine synthesis blockade did not impact the ability of bacteria to enter the Tel 4h post-injection (fig. S2, E to J). In response to injection of DFMA-DFMO treated bacteria, trout *odc* or *samdc* expression levels did not change in the Tel or Hyp at the time points tested (Fig. 3C and fig. S5, A to D). Combined, these results indicate that bacterial-derived polyamines selectively modulate host polyamine synthesis pathways in the Tel but not the Hyp in rainbow trout.

We next asked if changes in the expression of polyamine synthesis genes in the trout Tel correlated with changes in the Tel polyamine levels. We measured putrescine, spermidine and spermine levels in telencephalic tissue extracts under the different experimental conditions using ELISA. Systemic injection of gut commensals resulted in a 2-fold increase in telencephalic putrescine levels at both 4 hours and 1 day post-injection (Fig. 3G). Conversely, a sharp decrease in spermidine, and spermine levels was observed in commensal-injected trout, with a 3-5-fold reduction at both time points (Fig. 3H and I). Putrescine, spermidine and spermine levels in the Tel of trout injected with DFMA-DFMO treated bacteria were not different from vehicle controls. (Fig. 3G to I). These results indicate that bacteria-derived polyamines alter host polyamine metabolism in the Tel resulting in putrescine accumulation and downstream reduction of spermidine and spermine.

### Bacterial-derived polyamines selectively modulate telencephalic cytokine responses

Polyamines play crucial roles in innate immunity in mammals (66–68). We therefore asked if polyamines from microbiota could drive the expression of pyrogenic cytokines in the trout brain and hypothesized that this may occur in a region-specific manner. We administered DFMA-DFMO treated or untreated bacteria to animals (Fig. 4A) and measured *il-1b*, *tnfa*, and *il-6* transcript levels at 4 hours and 1 day post-injection. Untreated bacteria augmented the expression of *il-1b* and *tnfa* in the Tel 4 hpi. In sharp contrast, DFMA-DFMO treated bacteria failed to induce any significant changes in the expression levels of these cytokines (Fig. 4B and D). Additionally, the expression pattern of *il-6* in the Tel remained consistent across all treatment groups and the vehicle control (Fig. 4F). Conversely, the Hyp, which exhibited no changes in bacterial loads or in the expression of polyamine synthesis enzymes following injection, was agnostic to the ability of bacteria to synthesize polyamines. Bacteria injection, whether treated with DFMA-DFMO or untreated, resulted in an increase in the expression levels of *il-1b*, *tnfa*, and *il-6*, with fold changes ranging from 2 to 3, 4 hpi (Fig. 4C, E and G). Subsequently, the expression levels of these cytokines returned to their normal baseline within 1 day post-injection. These results suggest that polyamines derived from systemic bacteria and/or from the bacteria that penetrate the Tel directly modulate pyrogenic cytokine responses in the Tel but not the Hyp.

**Figure 4:**
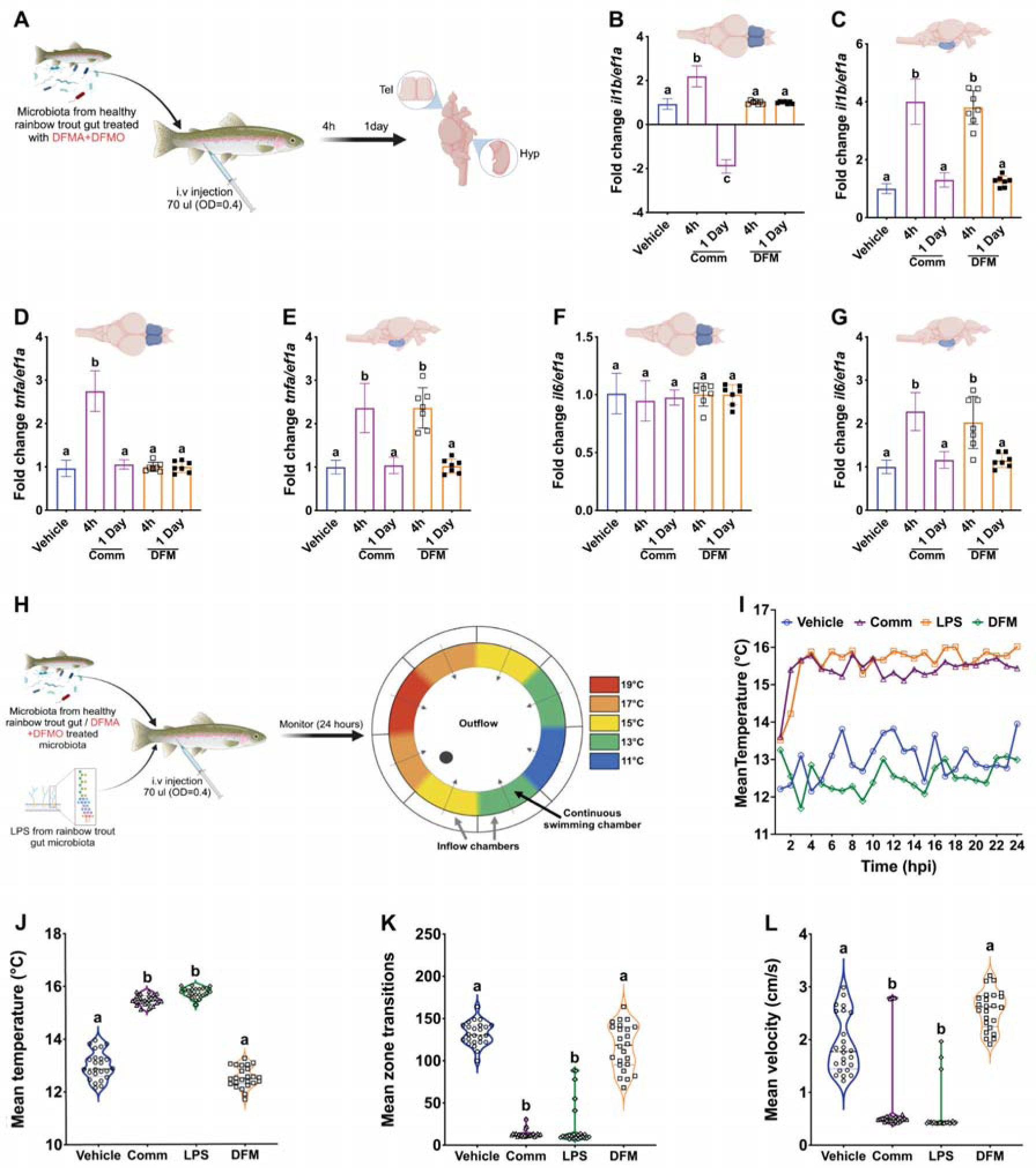
Systemic inflammation triggers pyrogenic cytokine expression and behavioral fever in a polyamine-dependent manner. (A) Schematic representation of the rainbow trout in vivo model used in this study. Trout received a single i.v injection of gut commensal bacteria treated or not with DFMA-DFMO to inhibit polyamine synthesis. (B-G) Changes in pro-inflammatory cytokine gene expression, *il-1b*, *tnfa*, and *il-6* in the Tel (B, D, F) and Hyp (C, E, G) 4 hours and 1 day post-injection of gut bacteria treated or not with polyamine synthesis inhibitors (N=7 fish/group). Expression values from the vehicle and commensal injected animals are the same as in Fig. 1. Statistical significance was tested at 95% interval using one-way ANOVA followed by Tukey’s post-hoc test (p<0.05). (H) Behavioral fever paradigm used in this study with a continuous ring-shaped swim chamber allowing rainbow trout to exhibit thermal preference between 11°C and 19°C. (I) Mean preferred temperature over the 24-hour observation period post-injection in vehicle, commensal injected, commensal + DFMA-DFMO (DFM) or LPS injected trout. (J) Mean preferred temperature of rainbow trout 4 hpi in each experimental group. (K) Mean zone transitions 4 hpi in each experimental group. (L) Mean swimming velocity 4 hpi in each experimental group. Different letters indicate statistically different groups by Brown-Forsythe and Welch’s one-way ANOVA test (p<0.05) (N=5).

### Systemic bacteria induce behavioral fever in rainbow trout in a polyamine-dependent manner

Behavioral fever is teleosts has been demonstrated in several models including viral and bacterial infections, killed bacterial challenges, and exposure to inflammatory agents like LPS or zymosan (30, 32, 33). However, whether the presence of microbiota in the systemic compartment induces behavioral fever in teleosts is currently unknown. We utilized a specialized annular thermal preference tank recently developed to document behavioral changes during fish fever (33) to assess the responses of four experimental groups (Fig. 4H). We observed that both i.v injection of gut commensals and LPS isolated from gut bacteria induced behavioral fever. Both groups reached a peak preferred temperature of 16°C (Fig. 4 I to L). However, fish injected i.v with gut commensals exhibited a more rapid onset of warm-seeking behavior compared to the LPS-injected group (Supplementary figure 4A). Two lethargy indicators were also evident in the commensal-injected and LPS-injected groups. Both groups displayed a marked decrease in swimming mobility and velocity (Fig. 4 K and L). Notably, maximum lethargy induced by systemic commensals occurred 2 hours earlier than by LPS (fig. S6, A and B). In contrast, treatment of bacteria with DFMA-DFMO prior to i.v injection prevented the increase in thermal preference and fever-associated lethargy behaviors. These fish showed no significant deviation from those behaviors in the vehicle-injected group (Fig. 4 I to L). Across all measured parameters, the response of animals injected with DFMA-DFMO treated commensals mirrored the behavioral patterns shown in the vehicle-injected group throughout the 24-hour study duration (Fig. 4 I, fig. S6, A and B). Together, these experiments revealed that 1) systemic inflammation induces behavioral fever in trout, and 2) commensal-derived polyamines are necessary to induce behavioral fever in trout under a scenario where systemic inflammation facilitates microbial translocation into the telencephalon.

## Discussion

Sickness behaviors and fever are intricate, coordinated neurobehavioral processes that enhance survival rates in vertebrates (12, 29, 69–72). As such, the vertebrate brain must sense and respond to peripheral and systemic insults to help the host return to homeostasis. The complex neuronal circuitry and immune mediators involved in mammalian sickness behaviors are fairly well characterized, reaching higher order brain regions beyond the Hyp (9–13). However, the neuroimmunological basis for sickness behaviors in ectotherms remains poorly understood.

Teleost fish, like other ectotherms, display sickness behaviors in response to infection characterized by reduced locomotor activity, social withdrawal, reduced appetite and behavioral fever (29, 32, 33, 35). Until now, the POV of the Hyp has been the only brain region implicated in responses during sickness behaviors and fever in teleosts (30–32, 73, 74). We hypothesized that higher order processing regions of the teleost brain, specifically the Tel, must participate in the neuroimmune response to systemic inflammation.

Here we present two novel acute systemic inflammation models in rainbow trout that mimic the presence of leaked microbiota or their products in the systemic compartment. Since teleost fish are highly refractory to *E. coli* LPS injections (47–49), we reasoned that direct injection of commensals or commensal-derived LPS i.v would be a more biologically relevant model of acute systemic inflammation than commercial LPS sources. In support, both *in vivo* models triggered canonical pyrogenic cytokine responses in the Hyp by 4 hours, as previously demonstrated by others using infection models (35, 40). In contrast, while *il-1b* and *tnfa* expression also increased in the Tel, *il-6* responses were not induced. In mammals, IL-6 can have pro-and anti-inflammatory roles in the brain (75–77). When injected systemically or into the brain, IL-6 causes some sickness behaviors in mammals (78), with unique biological activities compared to IL-1b (78). For instance, following a high dose of LPS injection, IL-1b knockout mice display reduced fever compared to controls, however, IL-6 knockout mice have normal fever responses (79). Additionally, IL-6 does not induce septic shock (80, 81) and injection of anti-IL-6 directly into the anterior POA suppresses PGE2 (82) and Cox2 (83). In teleosts, we know very little about the biological functions of IL-6. However, a study found that trout recombinant IL-6 reduces *il-1b* and *tnfa* expression in macrophages *in vitro* (84), suggesting that IL-6 responses dampen inflammation in trout. Assuming a similar function in the brain, *il-6* expression in the hypothalamus would reduce *il-1b* and *tnfa* expression creating a negative feedback loop that limits inflammation, whereas in the Tel, *il-1b* and *tnfa* effects should be prolonged. Intriguingly, trout have two fish-specific IL-6 family genes, the ciliary neurotrophic factor (CNTF-like) and M17 and CNTF-like is highly expressed in the brain (85). Thus, we cannot rule out that one or both molecules play alternative or complementary functions to canonical IL-6 in the teleost brain.

The BBB becomes leaky in response to systemic inflammation such as LPS injection, a phenomenon well-documented in mammals (50, 51). Loss of BBB integrity in response to LPS injection varies across different brain regions in mammals (50). Similarly, our studies indicate that the teleost BBB vulnerability to systemic inflammation varies among brain regions and that the Tel BBB is particularly susceptible to disruption in rainbow trout. Whether this is due to the unique cellular and molecular composition of the Tel’s BBB remains open for further investigation. Interestingly, our study surprisingly found that, in rainbow trout, circulatory bacteria transiently penetrate the Tel but not other brain regions under both acute (intravenous injection) and chronic (DSS-induced colitis) conditions. Prior research identified the insular cortex as a critical detector of peripheral inflammation in mice during DSS-induced colitis (16). However, this study did not explore whether bacterial translocation from the gastrointestinal tract to the bloodstream resulted in consequent bacterial infiltration into the brain. Nonetheless, the rapid and transient increase in bacterial loads observed in the trout Tel suggests a controlled mechanism by which the teleost brain may directly detect or sample changes in blood-borne bacteria. Whereas bacterial presence in the mammalian brain is thought to be a sign of disease (86–89), regulated bacterial incursions may not significantly harm the teleost brain. This hypothetical scenario is supported by the fact that, compared to mammals, the teleost brain harbors a higher number of immune cells under normal conditions (90) and possesses remarkable regenerative capabilities following injury (91–95).

We have recently reported that the trout brain has a resident bacterial community (55). Here we show that significant shifts in microbial community composition occur in the trout Tel following an i.v injection of gut bacteria. Following i.v injection, some bacteria will rapidly be eliminated by the blood immune system. Yet, we noted changes in the telencephalic microbial community 4 hpi that persisted by 1 day. We noted expansions of specific taxa such as Ruminococcaceae, Streptococcaceae and Mycoplasmataceae at 4 hours, that were mostly undetectable by 1 day indicating a transient colonization. This may be explained by rapid elimination of bacteria by the trout brain’s innate immune response or failure to outcompete other resident microbes. In contrast, Fusobacteriaceae and Chitinibacteraceae were expanded in the Tel after 1 day, potentially indicating longer-term dysbiosis in the brain following a single systemic inflammation event. We currently do not know how systemic bacteria penetrate the trout Tel, although our data suggest that the greater BBB disruption of the Tel may facilitate bacteria penetration. In support, we recently reported that mature Chinook salmon, where the gut barrier is disrupted, LPS levels increase in circulation and bacterial loads increase in the Tel (55). Thus, our findings suggest that teleosts may experience shifts in brain microbiomes throughout their lifespan, some associated with physiological and developmental changes, some because of stress, inflammation or infection.

Polyamines are polycationic compounds produced by eukaryotic and prokaryotic cells. In the vertebrate nervous system, polyamines are involved in neurodevelopment, synaptic plasticity, cognition, aging, and behavior (96–99). Importantly, polyamine metabolism and accumulation of putrescine are thought to be a protective biochemical program following brain insults in both mammals and fish (100–103). Our results indicate that specific regions of the trout brain (the Tel), but not others (the Hyp) quickly respond to systemic inflammatory cues by shifting polyamine metabolism. Specifically, we observed an accumulation of putrescine and a decrease in spermidine and spermine levels in the Tel. In support, putrescine was shown to accumulate in the goldfish brain following cold exposure suggesting that this protective response helps fish adapt to environmental changes (100). We also noted drastic decreases in telencephalic spermine levels. Since spermine inhibits proinflammatory cytokine responses, at least in humans (104), decreasing spermine levels in the Tel may be an effective mechanism to ensure pyrogen production, a hypothesis that merits further investigation. It is intriguing that changes in polyamine metabolism were not evident in the Hyp, reinforcing the notion that compartmentalized responses to systemic inflammation occur in the teleost brain, with the Tel being the focal point of polyamine-mediated responses.

The role of polyamines in mammalian fever responses remains unclear, with studies concluding either a lack of function as pyretics (105) or their antipyretic functions (106, 107). However, polyamines are precursors to gamma-aminobutyric acid (GABA) and GABA is known to act on the POA regulating body temperature by inhibiting heat production in mammals exposed to hot temperatures (108, 109). Interestingly, a recent study in human ulcerative colitis (UC) patients detected significant increases in serum N1-acetylspermidine levels, a bacterial-derived polyamine, in patients with moderate to severe UC compared to those with inactive disease (110) suggesting a role for N1-acetylspermidine in the pathology of UC or as a biomarker for disease severity. Interestingly, 40% of UC patients suffer from low fever of unknown sources (111) and therefore the possibility that bacterial-derived polyamines mediate this phenomenon is worth further investigation. A critical question of our study is whether the changes in polyamine metabolism detected in the Tel as well as subsequent behavioral fever responses require the incursion of microbiota into the Tel or they occur independently, for instance in response to the presence of bacteria in the systemic compartment Answering this question would require injecting bacteria mutants that lack the ability to enter the Tel so that we can disentangle systemic versus direct Tel responses.

## Conclusion

In conclusion, the present study suggests that polyamine metabolism ties immune regulation and behavioral adaptations to inflammation in teleosts, the Tel being the hub for this coordinated response. Yet, many questions remain to be answered, including the exact neuronal inhibitory and excitatory circuitry that connects the POV with the Tel during sickness behaviors and fever in ectotherms. Given that teleosts appear to live in symbiosis with bacteria during healthy states, as evidenced by their blood and brain microbiomes, we postulate that the Tel acts as a major brain sensor of systemic inflammatory states, allowing teleosts to rapidly respond to shifts in blood bacterial levels deviating from homeostasis.

## Materials and Methods

### Isolation of gut bacteria from juvenile rainbow trout

Rainbow trout, initially weighing 0.2 grams, were sourced from Trout Lodge (Washington, USA). They were maintained at a constant temperature of 16°C and fed daily with 2% of their body weight using commercial trout diet (Skretting). Feeding was discontinued one day prior to the collection of gut commensal bacteria. Trout (N=3 per collection group, mean body weight 23±3g) were euthanized using an overdose of MS-222 in water (Syndel, 200 ppm). Animals were then perfused by a cardiac injection of sterile PBS-heparin solution (Sigma Aldrich) to clear any blood from the gut-associated vasculature. Subsequently, the intestines were carefully removed and dissected lengthwise. Following mucus scraping, bacteria were isolated as previously described. The bacterial pellet was then washed in triplicate with a 1% PBS-BSA solution (pH 7.2), resuspended in sterile PBS and all five pellets were combined. The suspension was adjusted to achieve an optical density (OD) of 0.4 and injected using heparinized syringes.

### Inhibition of polyamine synthesis in isolated trout gut bacteria

To irreversibly suppress polyamine synthesis in isolated gut commensals from rainbow trout, we used the inhibitors DFMA (α-difluoromethylarginine) and DFMO (α-difluoromethylornithine) as previously described (65, 112, 113). Pilot experiments were performed to assess the growth kinetics of isolated gut bacteria in the presence of varying concentrations of each pharmacological agent tested independently (1 mM, 0.5 mM, 0.25 mM, 0.1 mM, 0.05 mM, 0.025 mM, 0.01 mM, and a control vehicle (PBS). Gut bacteria were adjusted to an OD of 0.05 in TSB and XX ul of the suspension were pipetted onto sterile flat-bottom 96-well plates (N=4/treatment). Plates were incubated at 30°C under continuous shaking. Optical density was measured every three hours for 24 hours in an iMark microplate reader (Bio-rad). Next, we evaluated the effects of growing gut bacteria with equimolar concentrations DFMA and DFMO concurrently using the same experimental set-up. These experiments identified the maximum concentration of DFMA and DFMO that could be used without impacting trout gut bacterial growth, which was 0.25 mM each.

The bacterial cultures were therefore incubated with a solution containing 0.25 mM of each DFMA and DFMO in phosphate-buffered saline (PBS) at a temperature of 4°C for a duration of 2 hours as previously described. Following the treatment period, the bacteria were then pelleted through centrifugation at 7,500 g for 20 minutes at 4°C. Next, the bacterial pellets underwent three separate washes with PBS. After each wash, the suspended bacterial cells were again centrifuged at 7,500 g for 20 minutes at 4°C to ensure the complete removal of any residual DFMA and DFMO. The optical density of the DFMA-DFMO treated bacterial suspension was adjusted to OD=0.4. This standardized bacterial preparation was then administered to the animals by i.v injection into the caudal vein.

### Lipopolysaccharide (LPS) Extraction

Isolated bacteria from rainbow trout gut were pellet down by centrifugation at 4°C for 25 min at 10000 g. LPS was isolated using the LPS Isolation Kit (MAK339, Millipore Sigma). In brief, the bacterial pellet underwent lysis via continuous pulse sonication for three 30-second intervals at 10 watts, utilizing a lysing buffer from the kit at a ratio of 1/10 mg/μL pellet to lysing buffer. Next, samples were incubated on ice for 10 minutes and subsequently centrifuged at 4°C for 10 minutes at 2,500 g. The supernatant was carefully transferred to a new 1.5 mL tube. Proteinase K was then added to achieve a final concentration of 0.1 mg/mL, and the mixture was incubated at 60°C for 60 minutes. Following incubation, the heated lysate was centrifuged at 4°C for 10 minutes at 2,500 g, and the supernatant was subjected to vacuum drying at 25°C. The dried sample was then reconstituted in an appropriate vehicle. The purity of the extracted LPS was evaluated using the total carbohydrate assay kit (MAK104, Millipore Sigma) which determined that the purity was >99.5%.

### In vivo systemic inflammation models

Juvenile trout (23 ±3.2 g body weight) were first sedated by MS-222 anesthesia in water (Syndel, 100 ppm). Animals (N=7/experimental group) were i.v injected with 70 μl of vehicle, isolated commensal bacteria (OD=0.4), LPS or DFMA-DFMO treated bacteria (OD=o.4) through the caudal vein. After 4 hours and 1 day, trout were euthanized by an overdose of 200 ppm MS-222 anesthesia (Syndel). Prior to tissue collection, animals were transcardiaclly perfused with sterile PBS-heparin solution (Sigma Aldrich) prior to tissue collection. The injections comprised 70 μl of either a vehicle, isolated commensal bacteria with OD=0.4, lipopolysaccharide (LPS), or bacteria treated with DFMA-DFMO at an OD of 0.4. Trout were euthanized at 4 hours and 1 day post-injection by an overdose of MS-222 in water until opercular movement stopped (Syndel). Prior to tissue collection, a sterile PBS-heparin solution (Sigma Aldrich) was used for cardiac perfusion to ensure the removal of blood and maintain tissue integrity.

To model chronic dysbiosis in trout, dextran sodium sulfate (DSS, MP Biomedicals) was employed to induce colitis-like symptoms following the protocol by Ding *et al.* (56). This involved administering a 3% DSS solution (by weight/volume in water) by oral gavage to the fish for a consecutive period of 10 days. Throughout the duration of the experiment, fish were fed daily 2% of their total body mass.

All animal procedures were approved by the University of New Mexico Institutional Animal Care and Use Committee under protocol number 22-201239-MC.

### Quantification of serum LPS levels

To assess the serum LPS levels, serum samples were first diluted at a 1:20 ratio using endotoxin free PBS (Millipore). Diluted serum samples were then subjected to a heat inactivation step, maintained at 70°C for 15 min. Following this, the endotoxin quantities in the serum were determined using the Pierce^TM^ chromogenic endotoxin quantification assay (Thermo Fisher Scientific) according to manufacturer’s instructions. To prevent any interference from pyrogenic substances, all laboratory consumables utilized in this process, such as pipette tips, Eppendorf tubes, and microtiter plates, were pre-validated to confirm their non-pyrogenicity.

### Quantitative real-time PCR for host gene expression profiling

Total RNA was extracted from the dissected tissues by the Trizol method. Tissue lysis was performed by the addition of two tungsten beads to 1.7 mL Eppendorf tubes containing the tissue and Trizol. The mixture was then subjected to vigorous shaking at 30 Hz for 5 minutes. Next, the homogenates/lysates were collected, and RNA extraction was performed following a standard chloroform/phenol extraction protocol, as delineated in previous studies. Total RNA was then quantified using a Nanodrop spectrophotometer. For cDNA synthesis, 1 μg of RNA underwent reverse transcription employing the Superscript III First Strand Synthesis System (Thermo Fisher Scientific, Catalog No. 18080051). Quantitative PCR (qPCR) was subsequently done using the SSOadvanced Universal SYBR Green Supermix (Bio-Rad, Catalog No. 1725270), with the annealing temperature set at 62°C using specific primers (Supplementary table 1) for the following genes: *il-1b, il-6, tnfa, odc1, odc2, samdc1, samdc2*. Changes in gene expression were quantified employing the Pfaffl method using *efa1* as the reference gene (114).

### DNA extraction and 16S rDNA library preparation

All the aseptic techniques were strictly practiced as previously described (55, 115). To lyse the specimens a pair of tungsten carbide beads was added to each tube, followed by vigorous lysis at 30 Hz for five minutes using the TissueLyser II (Qiagen). Post-mechanical lysis, we introduced 200 µl of a 1% CTAB (hexadecyltrimethylammonium bromide, Sigma) and 3 µl of proteinase K solution (100 mg/ml, Sigma) into each tube. The DNA extraction protocol was conducted in line with procedures previously established by Mitchell & Takacs-Vesbach (116). The purity and quantity of the extracted DNA were assessed using the Nanodrop ND 1000 spectrophotometer, with subsequent dilution of the DNA at either a 1:10 or 1:20 ratio. Our library construction for sequencing included both negative and positive controls, consistent with established norms within the microbiome research criteria (117, 118). To mitigate the influence of possible contaminants, we standardized the template DNA concentration for each PCR reaction at a minimum threshold of 200 ng. Despite the negative controls not meeting this concentration, the extraction elution was still subjected to PCR amplification. PCR was executed in triplicate for each biological specimen, with the resultant amplicons being combined before undergoing cleaning and library preparation. Every sequencing batch incorporated a positive control comprising DNA from seven laboratory-cultured bacterial strains to monitor sequencing accuracy. Additionally, a previously sequenced mouse colon DNA sample was used as a control in each sequence run to ensure methodological consistency.

To amplify the V1-V3 regions of the prokaryotic 16S rDNA, three distinct PCR reactions were performed on each sample using the designated primers 28F and 519R, as previously detailed (56). The resulting amplification mixtures were then pooled and subjected to a clean-up process by the AxyPrep Mag PCR Clean-up Kit (Thermo Fisher Scientific). The PCR products were then tagged using the Nextera XT Index Kit v2 Sets A to B (Illumina), with the amplicon concentrations normalized to 200 ng/µl via the Qubit high sensitivity dsDNA assay before pooling. An additional clean-up stage was done using the Axygen PCR clean-up system. The library was then sequenced on the Illumina NextSeq 2000 platform, by the 600-cycle reagent kit, at the Clinical and Translational Sciences Center, University of New Mexico Health Sciences Center.

### Bacterial load quantification in trout tissues

Bacterial loads within each sample were quantified via quantitative PCR targeting the 16S rDNA as previously described (55, 119). This involved the amplification of bacterial 16S rDNA sequences in triplicate, using primers specific to the V1-V3 hypervariable regions as established in prior research (56). The qPCR mixture was prepared in a 96-well format, containing 2 µl of standardized DNA/cDNA (10 ng/µl), a 2 µl aliquot of the primer mix, 6 µl of nuclease-free water, and 10 µl of SsoAdvanced Universal SYBR Green supermix (Bio-Rad). The amplification cycle commenced with an enzyme activation at 94°C for 1.5 minutes, followed by 33 cycles each consisting of denaturation at 94°C for 30 seconds, annealing at 52°C for 30 seconds, and extension at 72°C for 1.5 minutes. This was concluded by a final extension period at 72°C for 7 minutes and a subsequent cooling phase at 4°C. The process was carried out using the Bio-Rad CFX96 C1000 Touch Thermal Cycler. For quantification, the number of 16S rDNA copies in each sample was determined by comparison to a standard curve, which was generated using a range of E. coli 16S rDNA gene copies from 10^9^ down to 10 for the V1-V3 variable regions. All runs included negative controls with water as template.

### Microbiome data analyses

Microbial sequence data were analyzed using the Quantitative Insights Into Microbial Ecology 2 package (Qiime2, version 2023.7) (120). The Divisive Amplicon Denoising Algorithm 2 (DADA2) was utilized for partitioning demultiplexed reads into distinct amplicon sequence variants (ASVs) as described (121). These ASVs were then aligned against the Silva 16S rDNA reference database (version 138) for taxonomic characterization (122). Before proceeding to in-depth diversity analyses, samples were normalized to a uniform sampling depth of 4,000 reads per sample to ensure comprehensive rarefaction. The ensuing core diversity assessment was conducted while considering both temporal and experimental treatment factors. Using QIIME2, we computed various alpha diversity indices, including Shannon diversity, Chao1 richness, Pielou’s evenness index, and Faith’s phylogenetic diversity (PD). Beta diversity analyses (Weighted UniFrac distances) were performed in QIIME2 and principal coordinates analysis (PCoA) plots were visualized by the Qiime2R package in RStudio (version 1.3.959) (123). To infer the potential functional profiles of the microbial communities, we employed the PICRUSt2 (124) algorithm in Linux and visualized the outputs in RStudio (version 1.3.959) (123).

### Evan’s blue injection and extravasation quantification protocol

A 1% (w/v) stock solution of Evan’s blue dye was prepared in sterile PBS and subsequently filtered through 0.2 µm sterile filters to ensure purity. For the experimental procedure, 75 µL of the dye, either in the vehicle or mixed with the gut bacteria suspension, was injected into the caudal vein of rainbow trout, with each fish weighing approximately 10 g (± 1.35). The fish were then subjected to sampling at 4 hours and 1 day. Following euthanasia as above, each fish was perfused with a sterile perfusion solution directly through the heart to ensure the thorough removal of residual blood and the albumin-Evan’s blue complex from the vasculature. Prior to perfusion, the caudal vein and secondary branchial arch veins were incised to facilitate removal of any residual blood. Subsequently, the olfactory bulbs, telencephalon, hypothalamus, optic tectum and cerebellum were dissected. The weight of each dissected brain part was recorded, and the tissues were immediately immersed in 50% (w/v) trichloroacetic acid (TCA) and preserved on ice as described (52). Next, the tissues were subjected to a mechanical lysis step consisting of a 5-minute shaking at a frequency of 30 Hz. The resulting homogenate was then centrifuged at 10,000 rpm for 20 minutes at 4°C (52). The quantification of extravasated Evans Blue in each sample was achieved through colorimetry, comparing the absorbance of the samples to a standard curve generated from a known serial dilution of Evans Blue in 50% TCA at 610 nm (53). The calculated dye concentrations were then normalized to the weight unit of each sample, allowing for accurate comparative analysis between samples.

### Polyamine quantification by ELISA assay

Collected specimens were lysed using 1% Triton X-100 (v/v) (Sigma) as described (125). The polyamine profile of each sample including putrescine, spermidine, and spermine levels were quantified employing commercially available ELISA kits (MyBioSource) following the manufacturer’s instructions.

### Behavioral assays

Rainbow trout (40±7.6 g), were acquired from Smoky Farm Ltd. (Red Deer, Alberta, Canada). Upon arrival, the fish were accommodated in opaque, continuous flow-through aquaria maintained at temperatures of 10-12°C. These aquaria were equipped with constant aeration to ensure appropriate living conditions. A minimum acclimatization period of 14 days post-transport was held, during which the fish were subject to continuous monitoring. The dietary regimen consisted of floating pellets, administered once daily (2% of body weight) to provide the necessary nutrients.

Water quality was rigorously controlled, with dissolved oxygen levels maintained between 6.5-7.0 ppm and pH levels sustained within the range of 6.8 to 7.8. Prior to any experimental manipulation, fish were randomly selected, netted, and anesthetized using a 100 ppm tricaine methane sulfonate solution (Syndel), adhering to the ethical guidelines sanctioned by the Canadian Animal Care and Use Committees as well as the Science Animal Support Services. Cervical dislocation was employed as the method of sacrifice, ensuring adherence to ethical standards aimed at minimizing animal distress during experimental procedures.

All specimens were housed within the Aquatic Facility of the Department of Biological Sciences at the University of Alberta. The facility replicated natural light conditions, providing a photoperiod of 12 hours of light followed by 12 hours of darkness. Experimental interventions included intravenous injections of live commensal bacteria (OD = 0.4), isolated LPS (sourced from the same stock as utilized by the Salinas laboratory), or commensal bacteria treated with DMFA/DMFO, alongside a vehicle control (N=5). Each fish was then placed in an annular thermal preference tank (ATPT) for a 24-hour behavior analysis period as described (33).

Behavioral assessments were conducted using high-resolution positional tracking, facilitated by an infrared Panasonic CCTV color camera (model WV-CP620) equipped with a 2X variable focal lens (model WV-LZ61/2S) and a centrally positioned overhead lighting system. Video recording was achieved using Bandicam software (Bandicam Company, USA). The analysis of recorded videos was performed with Ethovision XT, Version 11 (Noldus, Wageningen, Netherlands), allowing for the quantification of behavioral parameters such as changes in thermal preference, swimming velocity, and the frequency of transitions between thermal zones as previously described (33).

### Statistical analyses

To assess the levels of endotoxin in blood serum among various groups, we initially conducted tests for normality using both the F-test and Bartlett’s test. Group differences were evaluated using either unpaired t-tests or one-way Analysis of Variance (ANOVA), with subsequent analysis through Tukey’s post hoc test for specific comparisons. To evaluate bacterial loads, indicated by 16S gene copy numbers in different tissues, data underwent checks for normal distribution as mentioned. To examine variations across tissues and between groups’ bacterial loads, we utilized both the Brown-Forsythe and Welch’s ANOVA tests; the same methodology was utilized to analyze the behavioral data. To analyze differences in Shannon Diversity indices, we first performed normality tests as mentioned earlier. Due to the non-parametric nature of these data, differences among groups were determined using the Kruskal-Wallis test. In instances of significant disparities, post hoc pairwise analysis was conducted, applying corrections for multiple comparisons. For visual interpretation of Weighted UniFrac distances, Principal Coordinates Analysis (PCoA) plots were used. Significance in clustering differences on these plots between groups was ascertained through multivariate dispersion analysis. All statistical evaluations were performed using GraphPad Prism (version 10.0.3) and RStudio (version 4.2.3), considering a P-value of less than 0.05 as statistically significant.

### Data availability

All microbiome sequencing data is available from NCBI under BioProject number PRJNA1079394.

## Acknowledgments

Authors thank Darrel Dinwiddie and Kurt Buchmann for support with the Illumina library sequencing and the University of New Mexico Center for Advanced Research Computing (CARC) for computing resources and data storage. Schematics were made in Biorender.

---

**fig. S1.**
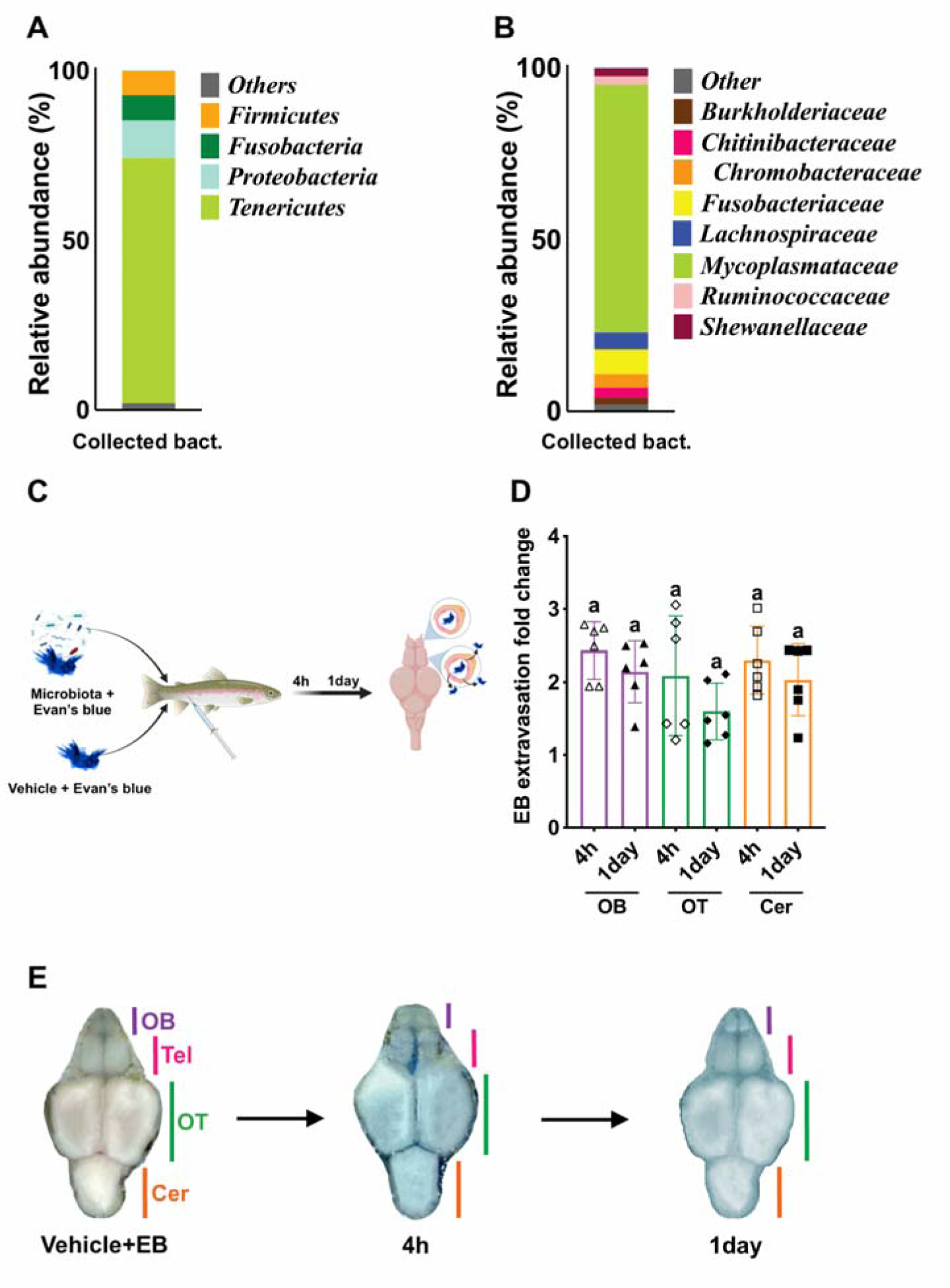
Blood-brain barrier integrity in various brain regions of rainbow trout during systemic inflammation. (A) 16S rDNA microbial community composition at the phylum (A) and family (B) levels of the isolated gut commensals prior to injection. (C) Schematic of the Evans blue extravasation assay. (D) Relative Evans Blue levels in the OB, OT and Cer 4 hours and 1 day post commensal injection (N= 6) relative to vehicle controls. (E) Images of whole dissected rainbow trout brains injected with vehicle or gut commensals at 4 hours and 1 day post-injection. Data were analyzed by one-way ANOVA followed by Tukey’s post hoc test (p<0.05).

**fig. S2.**
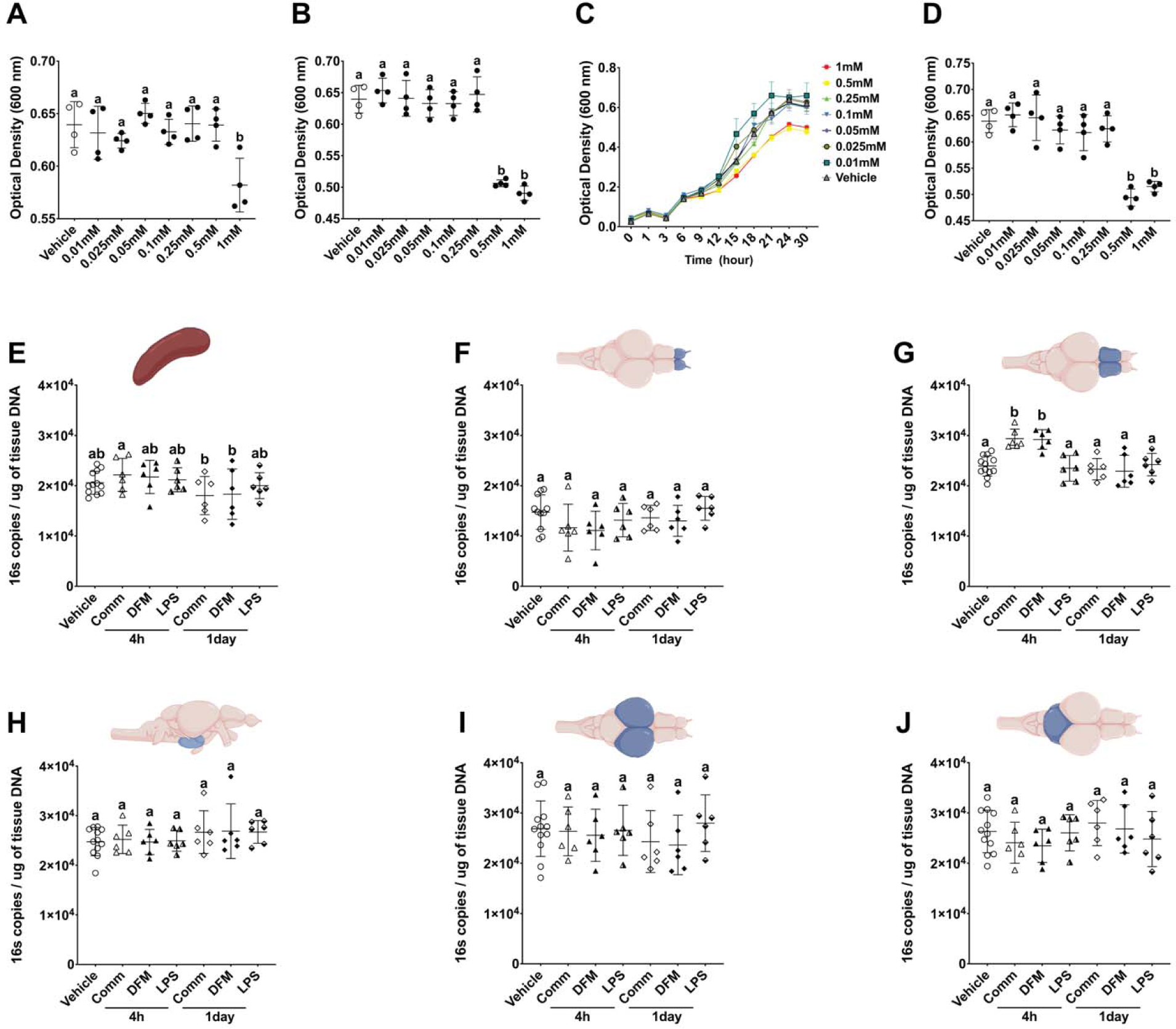
Impact of DFMA-DFMO on commensal bacteria growth and in vivo translocation of bacteria. (A) Optical density (O.D=600) of trout commensal bacterial cultures for 24 hours following incubation with different concentrations of DFMA (N=4). (B) Optical density of trout gut commensal cultures at the 24 hours endpoint following incubation with different concentrations of DFMO (N=4). (C) Growth curves of trout commensal bacteria exposed to combined DFMA-DFMO treatment for a 24 hour period (N=4). (D) Effect of combined DFMA-DFMO treatment on bacterial optical density at the 24 hours endpoint. Different letters indicate statistically significant groups by one-way ANOVA followed by Tukey’s post hoc test (N=4). (E-J) Quantitative PCR analysis of 16S rRNA gene copy numbers in the spleen, olfactory bulbs, telencephalon, hypothalamus, optic tectum, and cerebellum after i.v injection of commensal bacteria treated or not with DFMA-DFMO, or LPS at 4 hours and 1-day post-injection (N=6), The vehicle group has N=12 because it combines N=6 vehicle controls for each time point that show no significance differences between them. Statistical significance is denoted by distinct alphabetical markers as determined by ANOVA Welch’s test (p < 0.05).

**fig. S3:**
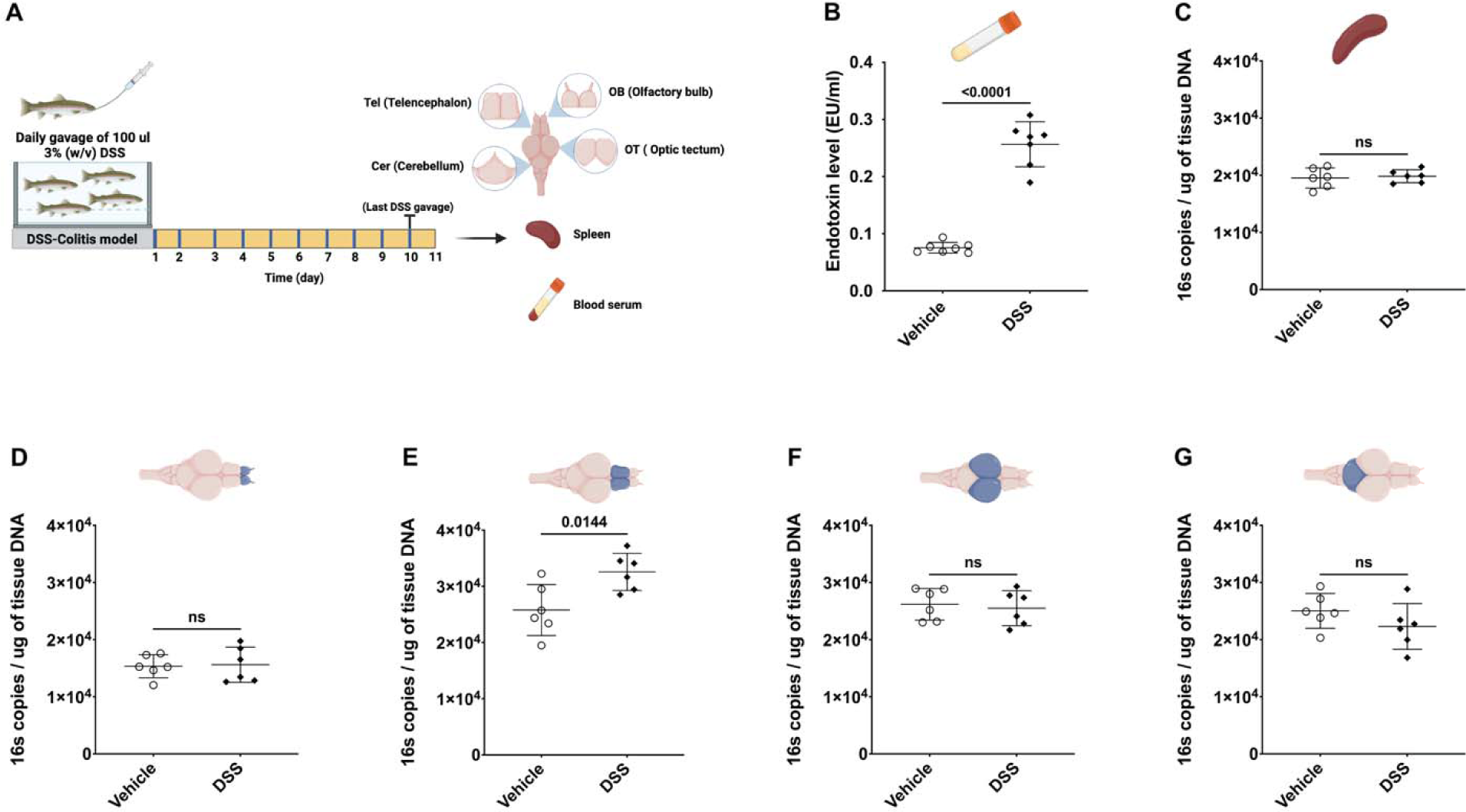
(A) Schematic illustration of the DSS-induced colitis model used in this study. (B) Comparative analysis of blood serum endotoxin levels between DSS treated animals and vehicle group. (C-G) Quantitative polymerase chain reaction (qPCR) assessment of 16S rRNA gene abundance in various brain regions and the spleen of both vehicle and DSS-induced colitis trout (N=6). Statistical significance is denoted as determined by unpaired two-tailed t-test (N=6).

**fig. S4.**
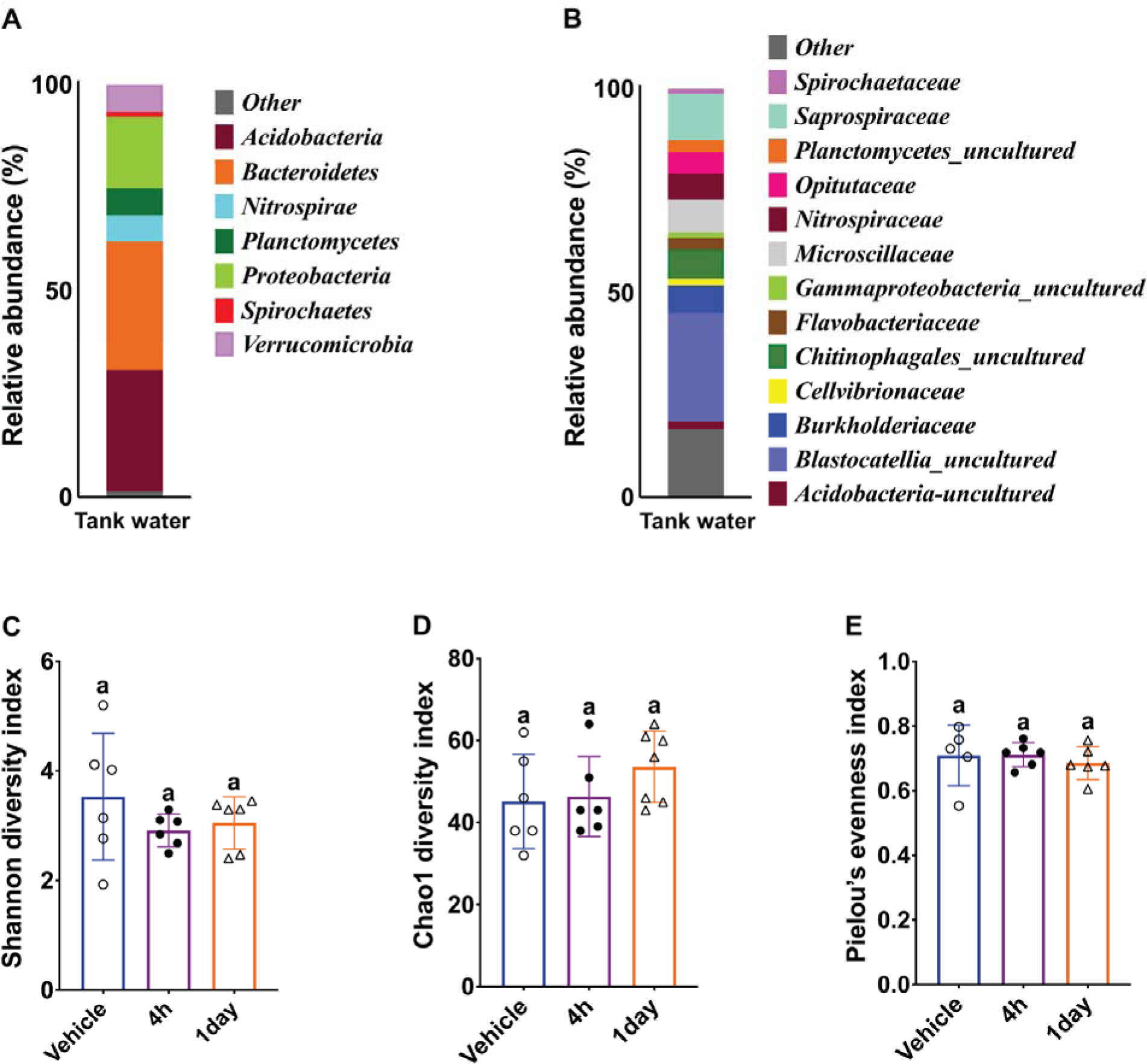
Microbial community analysis of tank water and rainbow trout telencephalon microbiome alpha diversity metrics post microbiota injection. Microbial community composition at the phylum (A) and family (B) levels of the tank water from the aquarium system where trout were maintained. (C-E) Alpha diversity metrics for the rainbow trout Tel microbial community at 4 hours and 1 day post gut commensal injection. Shannon diversity index (C), the Chao1 diversity index (D), and Pielou’s evenness index (E) (N=6). Different letters denote statistically significant groups (P<0.05) by Kruskal-Wallis test.

**fig. S5:**
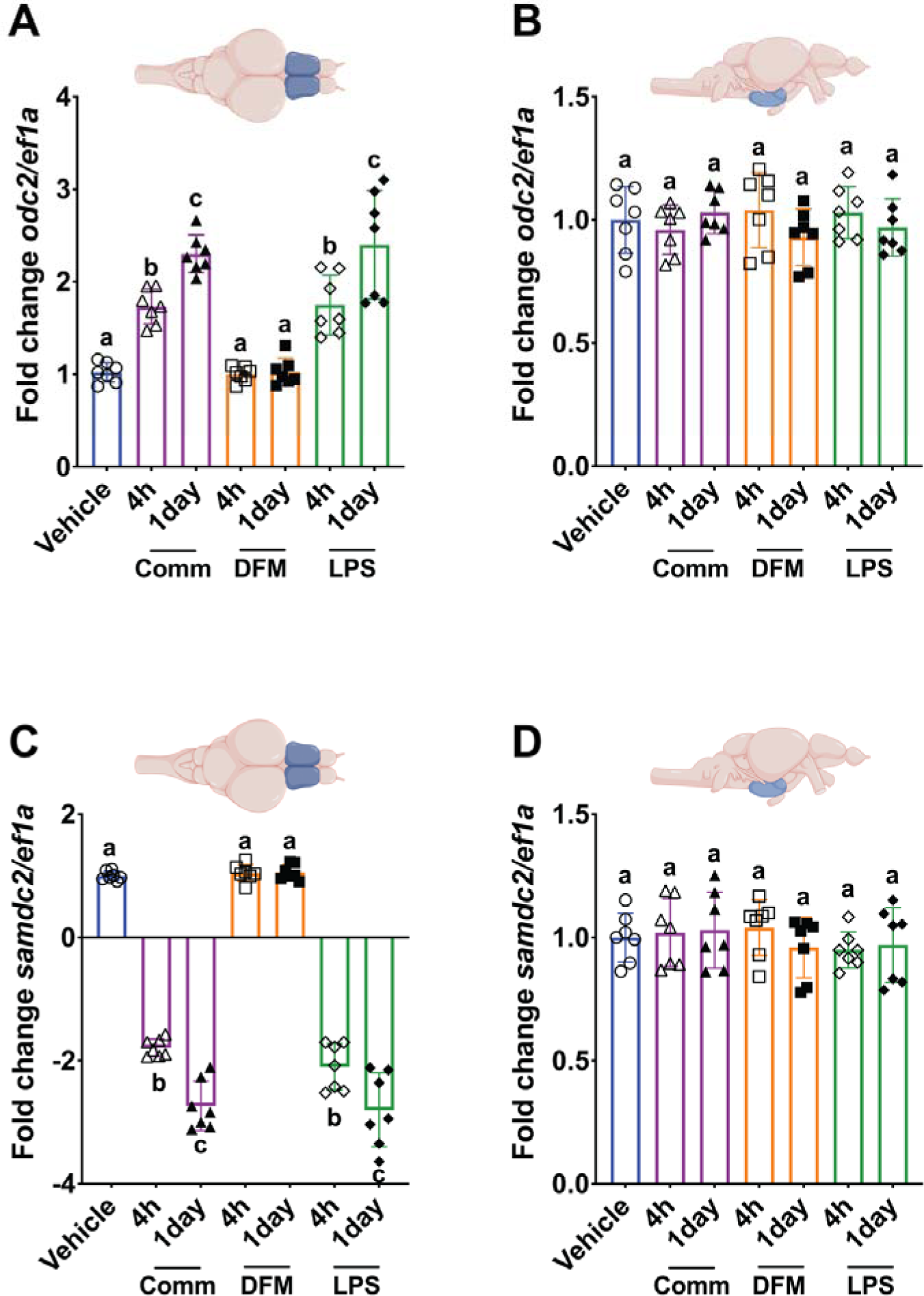
Changes in expression levels of the polyamine synthesis enzymes *odc2* and *samdc2* in the telencephalon and hypothalamus post-injection of gut commensals treated or not with DFMA and DFMO or LPS extracted from gut commensals as measured by RT-qPCR (N=6-7). Expression levels were normalized to *ef1a*. Data were analyzed used the Pfaffl method. Different letters indicate statistically significant groups based on one-way ANOVA followed by Tukey’s post hoc test (p<0.05).

**fig. S6.**
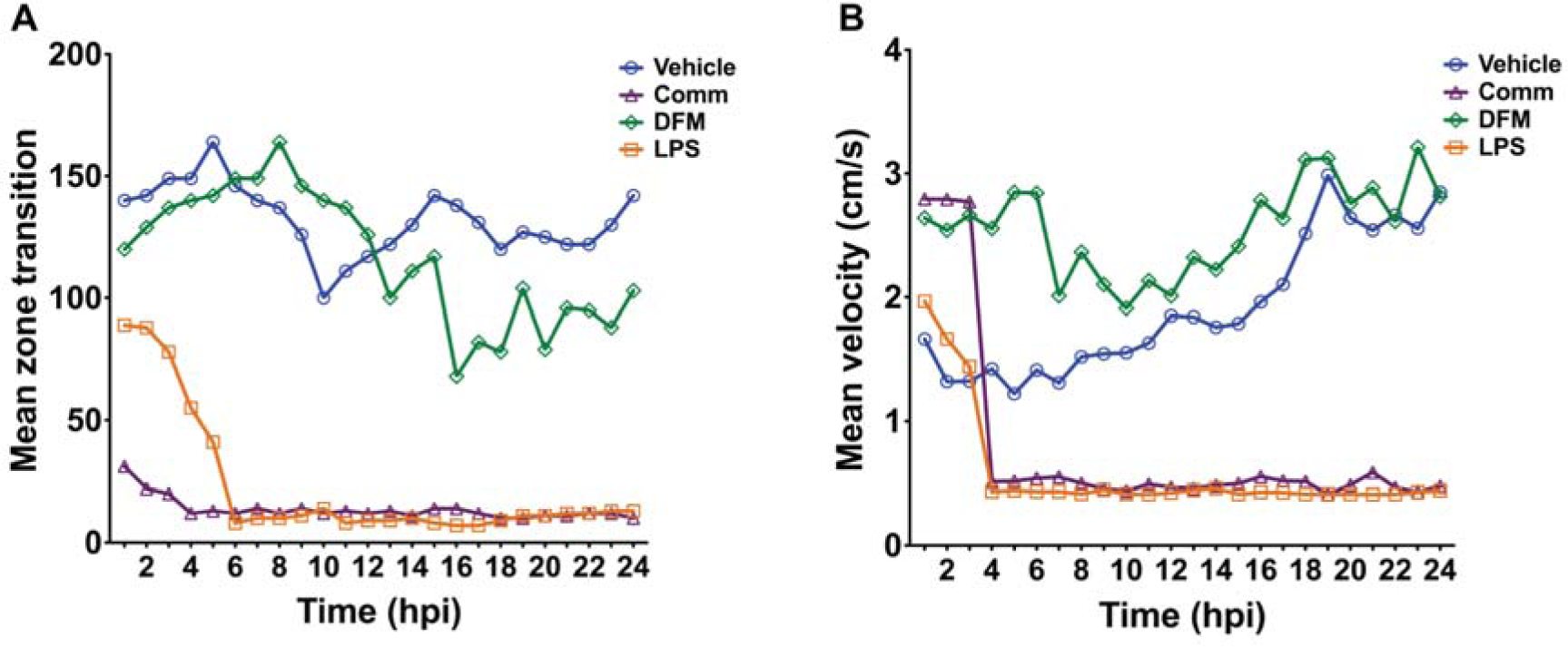
Behavioral responses in rainbow trout following microbiota and LPS Injections. (A) Mean zone transition frequency over a 24-hour period post-injection, reflecting the locomotor activity of rainbow trout in response to various treatments. The graph illustrates the trout’s exploratory behavior, with notable differences among groups treated with vehicle, gut commensals (Comm), DFMA-DFMO treated commensals (DFM), and LPS. (B) Mean swimming velocity over 24 hours post-treatment in the four experimental groups (N=5 fish/treatment).

**Table S1.**
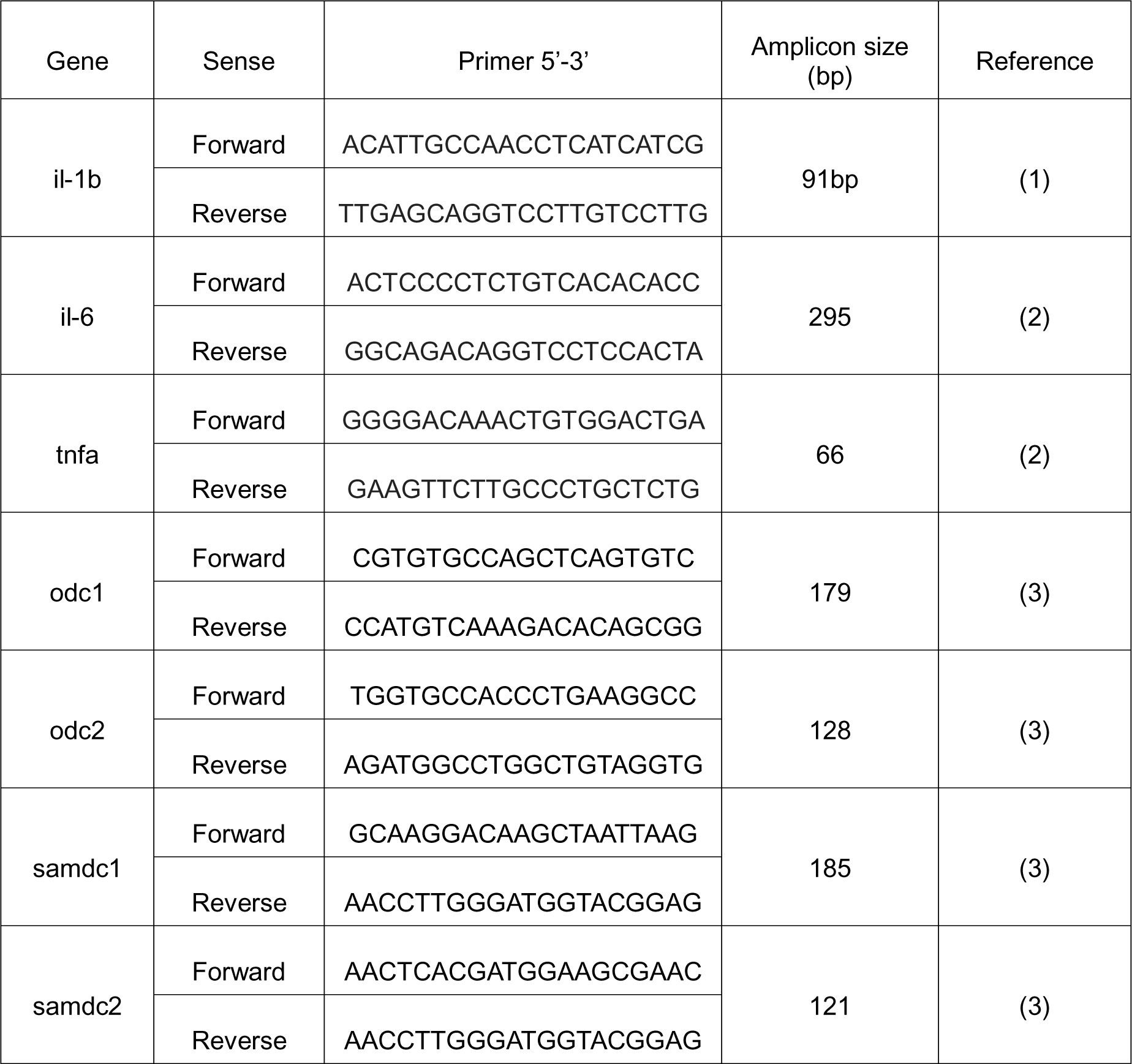
Primer sequences employed in this study, adapted from published sources.

